# Early apelin receptor activation attenuates elastase-induced emphysema and preserves endothelial apelin receptor signaling in mice

**DOI:** 10.64898/2026.05.12.724387

**Authors:** Tomoki Kishimoto, Ryunosuke Nakashima, Keisuke Kawano, Mai Uemura, Kota Nakajima, Noriki Takahashi, Choyo Ogasawara, Yukio Fujiwara, Mary Ann Suico, Hirofumi Kai, Tsuyoshi Shuto

## Abstract

Alveolar capillary endothelial cells are positioned adjacent to the alveolar epithelium and contribute to lung homeostasis and injury responses. Single-cell studies have identified aerocyte capillary endothelial cells (aCap), which are specialized for gas exchange, and general capillary endothelial cells (gCap), which contribute to endothelial maintenance and inflammatory signaling. Apelin and its receptor are differentially enriched across these endothelial compartments, but their roles in emphysema development remain incompletely understood. Using an elastase-induced emphysema model in male C57BL/6J mice, we combined bulk RNA sequencing, CIBERSORTx-based cell-type deconvolution, histology, inflammatory assays, pulmonary function testing, and pharmacologic activation of the apelin receptor with [Pyr^1^]-Apelin-13. At 24 hours after elastase exposure, the inferred fraction of gCap was reduced, and lung expression of apelin and the apelin receptor was decreased. Early [Pyr^1^]-Apelin-13 administration reduced lung inflammatory mediator expression, Ly6G-positive neutrophil accumulation, bronchoalveolar lavage neutrophil counts, and matrix metalloproteinase-9 activity. Early treatment also attenuated subsequent airspace enlargement, whereas treatment initiated after emphysema was established did not improve physiological or histological outcomes. In a chronic βENaC-transgenic mouse model, the inferred gCap fraction was maintained, the aCap fraction was reduced, and apelin receptor activation did not improve disease phenotypes. These findings suggest that early activation of the apelin receptor modifies acute inflammatory and endothelium-associated responses following elastase injury and limits emphysematous remodeling in mice. Together, these results support a time-sensitive role for apelin-APJ signaling during the early phase of emphysema development.

## Introduction

The distal lung is composed of specialized epithelial, endothelial, mesenchymal, and immune cell populations that cooperate to maintain the alveolar gas-exchange surface. Type I alveolar epithelial cells (AT1) cover most of the alveolar surface area, whereas type II alveolar epithelial cells (AT2) act as progenitor cells and contribute to surfactant production [1,2]. Alveolar capillary endothelial cells are positioned next to the alveolar epithelium, where they participate in alveolar-capillary barrier integrity and provide signals that support alveolar homeostasis.

Endothelial-epithelial crosstalk is increasingly recognized as an important determinant of alveolar injury and repair. Lung endothelial cells produce angiocrine factors, including thrombospondin 1, matrix metalloproteinase 14, and hepatocyte growth factor, that can influence epithelial progenitor behavior and alveolar regeneration [3–7]. Conversely, endothelial cell injury or apoptosis has been implicated in chronic obstructive pulmonary disease and experimental emphysema [8–11]. These observations suggest that endothelial dysfunction is not merely a consequence of alveolar injury but may contribute to disease progression.

Single-cell RNA sequencing studies have further refined the concept of the alveolar endothelium by identifying distinct capillary endothelial subtypes, including aerocyte capillary endothelial cells (aCap) and general capillary endothelial cells (gCap) [12,13]. aCap is specialized for gas exchange and constitutes a major component of the air-blood barrier, whereas gCap is involved in endothelial maintenance, repair, and inflammatory signaling. Human single-cell analyses of chronic obstructive pulmonary disease (COPD) have also reported alterations in the alveolar niche, including changes in endothelial programs and injury-associated states [14]. These findings raise the possibility that an imbalance in endothelial subtypes may influence the development of emphysema.

Apelin and its G protein-coupled receptor, commonly referred to as APJ, are candidate mediators of endothelial homeostasis. Apelin is enriched in aCap, whereas the apelin receptor is enriched in gCap [12,13]. The apelin-apelin receptor axis regulates angiogenesis, endothelial survival, inflammation, and vascular permeability in several contexts [15–19]. In lung injury models, apelin-related signaling has been reported to improve epithelial barrier integrity, endothelial barrier function, and inflammatory responses [20–23]. However, whether activation of the apelin receptor modifies early endothelium-associated responses during emphysema development remains unclear.

Based on these observations, we investigated whether apelin-APJ signaling is altered during emphysema development after acute epithelial injury and whether pharmacologic activation of APJ modifies the subsequent inflammatory and structural responses. We analyzed endothelial subtype-associated transcriptomic changes in elastase-treated mouse lungs and examined the effects of early and delayed [Pyr^1^]-Apelin-13 administration on acute inflammation, pulmonary function, and emphysematous remodeling. We also compared these findings with a chronic βENaC-transgenic model to evaluate the context dependence of apelin receptor activation.

## Materials and Methods

### Animals and ethics statement

Male C57BL/6J wild-type mice were obtained from Charles River Laboratories Japan (Atsugi, Japan). C57BL/6J-βENaC transgenic mice were generated and maintained at the Center for Animal Resources and Development, Kumamoto University, as previously described [24]. Mice were housed under specific pathogen-free conditions on a 12-hour light/dark cycle at 22–25°C with ad libitum access to chow and water. All animal experiments were approved by the Animal Care and Use Committee of Kumamoto University (Approval No. A2024-052) and were performed in accordance with institutional and national guidelines and ARRIVE recommendations. Mice were anesthetized by intraperitoneal injection of medetomidine (0.75 mg/kg; Nippon Zenyaku Kogyo, cat. no. 14111), midazolam (4 mg/kg; Sandoz, cat. no. 1124401A1060), and butorphanol (5 mg/kg; Meiji Seika Pharma, cat. no. 49872222889575). After a surgical plane of anesthesia was confirmed by loss of the pedal withdrawal reflex, blood was collected by cardiac puncture, and euthanasia was completed by exsanguination under deep anesthesia. Death was confirmed by cessation of respiration and heartbeat before tissue collection. Animals were monitored throughout the experiments, and all efforts were made to minimize suffering.

### Elastase-induced emphysema model and apelin receptor activation

For the elastase-induced emphysema model, 11-week-old male C57BL/6J mice received intratracheal administration of porcine pancreatic elastase (E0127; Sigma-Aldrich, Tokyo, Japan) at 2 mg/kg, dissolved in saline to 1 mg/mL [25]. For acute-phase analyses, [Pyr^1^]-Apelin-13 was administered intraperitoneally at 0.5 μmol/kg beginning 1 hour after elastase exposure, with an additional dose at 24 hours as required by the experimental schedule. Samples for acute-phase analyses were collected 24 hours after elastase exposure. For prevention experiments, [Pyr^1^]-Apelin-13 treatment was initiated 1 hour after elastase exposure and continued once daily for 3 weeks. For delayed-treatment experiments, once-daily [Pyr^1^]-Apelin-13 treatment was initiated 3 weeks after elastase exposure, when emphysema was already established, and continued until analysis. βENaC-transgenic mice were used as a model of chronic airway inflammation and mucus obstruction and were analyzed for endothelial subtype-associated signatures and responses to apelin receptor stimulation.

### Pulmonary function testing

Pulmonary function was measured using the flexiVent system (SCIREQ, Montreal, Canada) according to the manufacturer’s protocol. Anesthesia was induced as described above, and an 18-gauge needle was inserted into the trachea, and the mice were mechanically ventilated at 150 breaths/min. Inspiratory capacity (IC) was measured using a deep-inflation maneuver. Compliance and elastance were assessed using the forced oscillation technique. Forced expiratory volume in 0.1 second (FEV_0.1_), forced vital capacity (FVC), and the FEV_0.1_/FVC ratio were measured using a negative-pressure reservoir connected to the flexiVent system and analyzed with flexiVent software.

### Tissue preparation and histology

At the designated endpoints, the trachea and whole lungs were excised and washed with phosphate-buffered saline (PBS). For histological analyses, the left lung was immersed in 10% neutral buffered formalin and fixed at room temperature for 12–24 hours. After fixation, the left lung was divided into three parts and washed three times with PBS. Tissues were dehydrated through graded ethanol and processed for paraffin embedding by K.I. Stainer Co., Ltd. Paraffin-embedded lung tissues were sectioned for histological and immunohistochemical analyses. Tissue sections were stained with Alcian blue (pH 2.5), periodic acid-Schiff, and hematoxylin. Images were acquired using a BZ-X710 microscope (Keyence, Osaka, Japan).

### Morphometric analysis

Emphysema severity was quantified using the mean linear intercept (MLI), as previously described [25]. Briefly, representative lung fields were analyzed from Alcian blue/periodic acid-Schiff-stained sections, and the MLI was calculated as an index of airspace enlargement.

### Bronchoalveolar lavage fluid analysis

Bronchoalveolar lavage fluid (BALF) was mixed with Türk’s solution, and total leukocytes were counted using a Bürker-Türk hemocytometer. Samples were centrifuged at 800 × g for 10 minutes. Supernatants were collected for enzyme-linked immunosorbent assays and activity assays. Cell pellets were resuspended in RPMI1640 containing fetal bovine serum, cytospun onto glass slides, and stained with Diff-Quik for differential cell counting based on cellular morphology. Morphology-based BALF differentials were integrated with Ly6G immunohistochemistry to assess neutrophilic inflammation.

### Immunohistochemistry and quantification

Paraffin-embedded lung sections were deparaffinized with xylene and rehydrated through graded ethanol. Antigen retrieval was performed using Target Retrieval Solution. Endogenous peroxidase activity was quenched by incubation with 3% H_2_O_2_ in PBS or H_2_O_2_ in methanol for 15 minutes. After washing with PBS, sections were incubated overnight at 4°C with primary antibodies diluted in Antibody Diluent (Dako). The following primary antibodies were used: rabbit anti-Iba1 (019-19741; FUJIFILM Wako, Osaka, Japan) and rabbit anti-Ly6G (ab238132; Abcam, Cambridge, UK). After three washes with PBS, sections were incubated with Histofine Simple Stain Mouse MAX-PO(R) (Nichirei, Tokyo, Japan) for 1 hour at room temperature. Signals were developed using DAB substrate (425011; Nichirei), and sections were counterstained with hematoxylin. Whole-slide images were acquired using a NanoZoomer S20 Digital Slide Scanner (Hamamatsu Photonics, Hamamatsu, Japan).

Immunohistochemical staining was quantified using HALO image analysis software (Indica Labs, Albuquerque, NM, USA). Whole-slide images were imported into HALO, and representative color tones for hematoxylin-positive nuclei and DAB-positive staining were registered for each slide. Threshold settings were optimized using the real-time tuning function, and the entire slide was analyzed after confirming accurate recognition of DAB-positive staining. The percentage of DAB-positive cells among total cells was calculated for each marker.

### Immunofluorescence

Paraffin-embedded lung sections prepared as described above were deparaffinized and subjected to antigen retrieval in Target Retrieval Solution (Agilent Dako, Santa Clara, CA, USA) at 95°C for 20 minutes. Primary antibodies were anti-APJ (20341-1-AP; Proteintech, Rosemont, IL, USA) and anti-CD31 (sc-376764; Santa Cruz Biotechnology, Dallas, TX, USA). Secondary antibodies were Alexa Fluor 546-conjugated goat anti-rabbit IgG (A11010; Thermo Fisher Scientific, Waltham, MA, USA) and Alexa Fluor 488-conjugated goat anti-rat IgG (A11006; Thermo Fisher Scientific). Nuclei were counterstained with DAPI (Dojindo, Kumamoto, Japan). Images were acquired using a BZ-X710 microscope. APJ/CD31 double-positive signals were quantified as a histological index of APJ-positive endothelial cells, without assigning these cells to a specific endothelial subtype.

### RNA isolation and quantitative reverse transcription-polymerase chain reaction (qRT-PCR)

Total RNA was extracted from mouse lung tissues using RNAiso Plus (Takara Bio, Shiga, Japan). Reverse transcription was performed using PrimeScript RT Master Mix (Takara Bio) according to the manufacturer’s instructions. qRT-PCR was performed using TB Green Premix Ex Taq II (Takara Bio) and a CFX Connect real-time polymerase chain reaction system (Bio-Rad, Hercules, CA). Cycling conditions were 95°C for 3 minutes followed by 40 cycles of 95°C for 10 seconds and 65°C for 1 minute. Expression levels were normalized to 18S ribosomal RNA (rRNA) levels. Primer sequences are listed in Table 1.

**Table 1.**
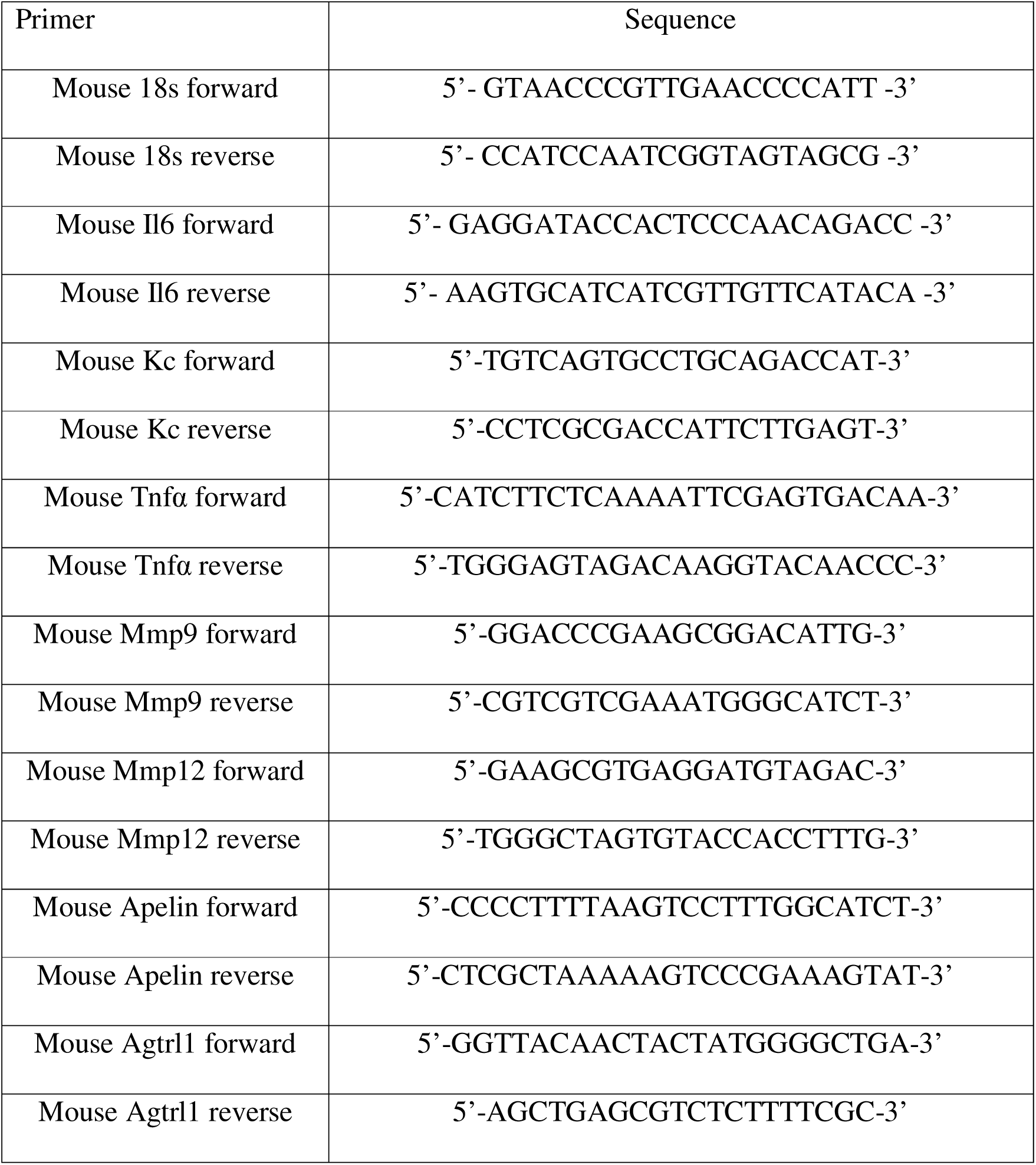
Sequences of primers for quantitative RT-PCR.

### Bulk RNA sequencing and CIBERSORTx analysis

Total RNA was extracted from mouse lung tissue using the Maxwell RSC simplyRNA Tissue Kit (Promega, Madison, WI). RNA concentration was measured with a NanoDrop One spectrophotometer (Thermo Fisher Scientific), and RNA integrity was assessed using the TapeStation 4200 system (Agilent Technologies, Santa Clara, CA). Polyadenylated RNA was isolated using the NEBNext Poly(A) mRNA Magnetic Isolation Module (New England BioLabs, Ipswich, MA). Libraries were prepared using the NEBNext Ultra II Directional RNA Library Prep Kit for Illumina, including RNA fragmentation, first- and second-strand complementary DNA synthesis, adapter ligation, and barcode indexing. Library preparation, quality control, and sequencing were performed at the Gene Research Center, Yamaguchi University, using a NovaSeq 6000 system.

Base calling and FASTQ conversion were performed using bcl2fastq. Reads were trimmed in CLC Genomics Workbench (QIAGEN, Hilden, Germany) and mapped to the Mus musculus GRCm39 reference genome from Ensembl. Gene expression was quantified as transcripts per million. Principal component analysis and Gene Ontology analysis were performed using RNAseqChef. Differentially expressed genes were identified using thresholds of |log[ fold change| > 1 and a false discovery rate < 0.05.

Cell-type-associated transcriptional composition was estimated using CIBERSORTx [26] with a custom signature matrix derived from the publicly available mouse lung single-cell RNA-sequencing dataset GSE168299. CIBERSORTx was run in relative mode using the default permutation setting. Batch correction was not applied before deconvolution; therefore, the resulting values were analyzed as inferred cell-type-associated fractions and interpreted as exploratory estimates, accounting for potential influences of sequencing batch, tissue composition, inflammatory transcriptional changes, and reference-matrix dependence. Bulk RNA-sequencing data from the elastase model are available under accession number GSE324765. Bulk RNA-sequencing data from C57BL/6J-βENaC-transgenic mice are available under accession number GSE300375 [27].

### Enzyme-linked immunosorbent assays (ELISA) and activity assays

Apelin was measured using a Mouse Apelin-13 ELISA Kit (LS-F28139; LifeSpan BioSciences, Seattle, WA). Keratinocyte-derived chemokine/CXCL1 was quantified using a Mouse CXCL1/KC DuoSet ELISA Kit (R&D Systems, Minneapolis, MN). Matrix metalloproteinase-9 (MMP-9) activity was measured using an MMP9 Activity Assay Kit (QZBMMP9M; QuickZyme Biosciences, Leiden, the Netherlands). Assays were performed according to the manufacturer’s instructions using bronchoalveolar lavage fluid, serum, or lung tissue lysates, as appropriate.

### Immunoblotting

Lung tissue lysates were analyzed by immunoblotting for MMP-9, with vinculin used as a loading control. Band intensity was quantified from digital images. Because MMP-9 can be produced by multiple cell types in injured lung tissue, immunoblot and activity data were interpreted as tissue- or bronchoalveolar lavage-level measurements rather than as evidence of a specific cellular source.

## Statistical analysis

Data are presented as mean ± SEM. Each biological replicate represents one mouse. Sample sizes were based on previous studies using elastase-induced and βENaC-transgenic mouse models, as well as feasibility considerations for animal experiments; no formal power calculation was performed. Animals were allocated to treatment groups before sample collection, and histological quantification was performed using digital image analysis. No data were excluded unless a technical failure occurred during sample processing or measurement. Comparisons among three or more groups were performed using one-way ANOVA followed by Dunnett’s multiple-comparisons test. Comparisons between two groups were performed using a two-tailed unpaired Student’s *t*-test. Statistical analyses were performed using GraphPad Prism 9.0 (GraphPad Software, San Diego, CA). A *P* value less than 0.05 was considered statistically significant.

## RESULTS

### Elastase injury is associated with early reduction of an inferred general capillary endothelial signature and decreased apelin receptor pathway expression

To examine endothelial-associated changes during emphysema development, we used an elastase-induced mouse model in which intratracheal elastase administration produces progressive emphysematous airspace enlargement over approximately 2–3 weeks [25,28]. Lung tissues were collected from control mice and from elastase-treated mice at day 1, week 1, and week 2 for bulk RNA sequencing (Fig 1A and 1B).

**Fig 1.**
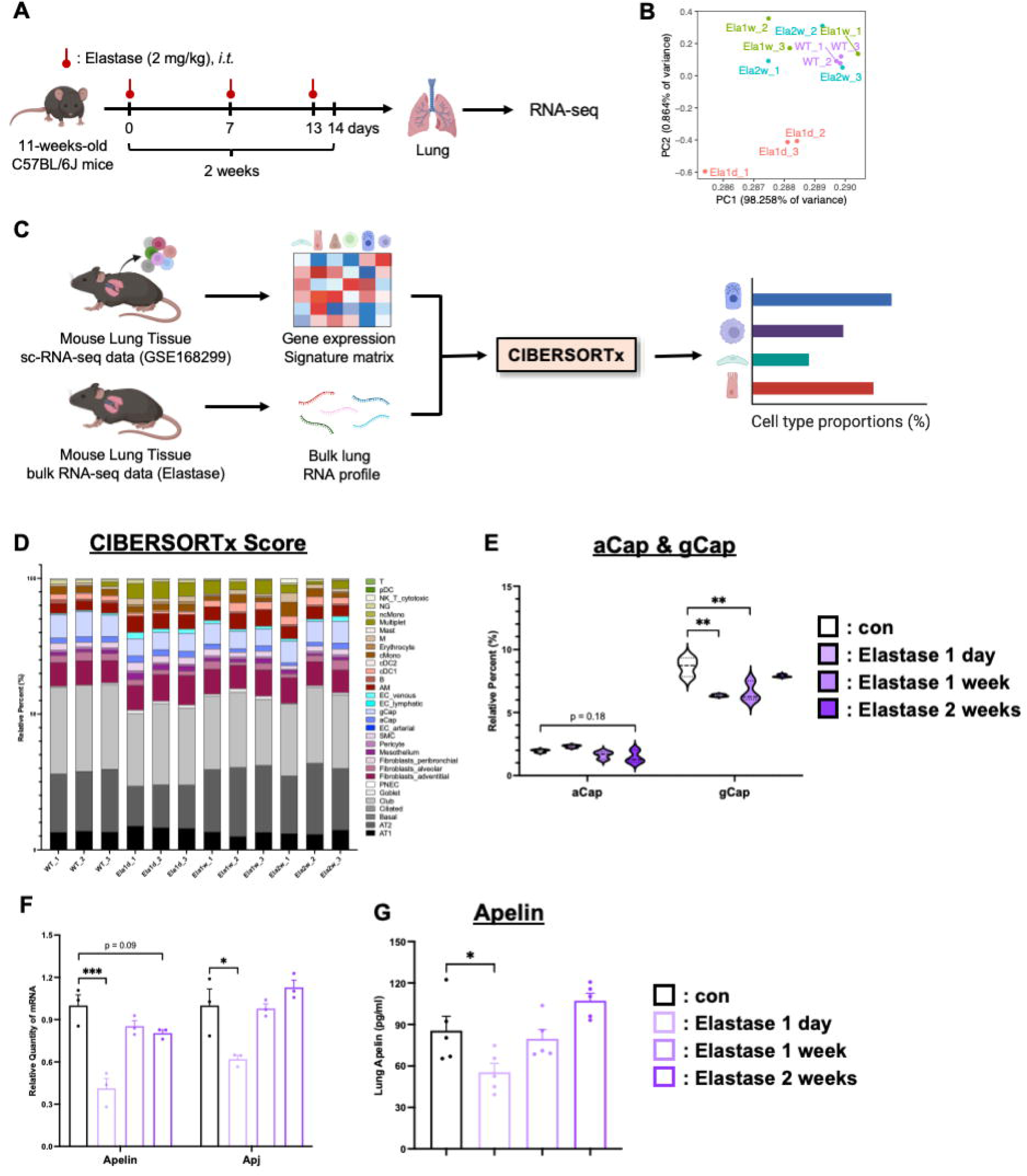
Elastase injury is associated with early endothelial subtype-specific transcriptional changes and reduced expression of the apelin receptor pathway. (A) Experimental design for elastase administration and lung RNA sequencing. Eleven-week-old male C57BL/6J mice received intratracheal elastase at 2 mg/kg, and lungs were collected at day 1, week 1, and week 2. (B) Principal component analysis of lung bulk RNA-sequencing data from control and elastase-treated mice. (C) Overview of the CIBERSORTx workflow. A signature matrix generated from mouse lung single-cell RNA-sequencing data (GSE168299) was applied to bulk lung RNA-sequencing profiles from control and elastase-treated mice. (D) Estimated proportions of lung cell-type-associated signatures inferred by CIBERSORTx. (E) Inferred aerocyte capillary endothelial (aCap) and general capillary endothelial (gCap) fractions. (F) Lung expression of *Apln* and apelin receptor/APJ-associated transcript in bulk RNA-sequencing data. (G) Lung apelin protein levels measured by enzyme-linked immunosorbent assay. Data are means ± SEM; n = 3–6 mice per group. *P* values were determined by one-way ANOVA with Dunnett’s multiple-comparisons test. \**P* < 0.05, \*\**P* < 0.01, \*\*\**P* < 0.001.

Gene Ontology analysis showed time-dependent transcriptional changes after elastase exposure (S1 Fig). At day 1, upregulated pathways were dominated by leukocyte chemotaxis, granulocyte migration, cytokine production, and related inflammatory processes. At week 1, cell cycle and tissue remodeling-associated terms were more prominent. At week 2, immune-associated pathways, including B cell receptor signaling and complement activation, were detected. These results are consistent with a transition from acute inflammation to later remodeling and immune responses.

We next estimated cell-type-associated transcriptional composition using CIBERSORTx with a mouse lung single-cell RNA-sequencing reference dataset (GSE168299) (Fig 1C). This analysis suggested that the inferred fraction of general capillary endothelial cells (gCap) was reduced after elastase exposure, most prominently during the early phase, whereas changes in aerocyte capillary endothelial cells (aCap) were less marked (Fig 1D and 1E). Deconvolution also suggested changes in epithelial, mesenchymal, and immune cell-associated fractions, including reduced type II alveolar epithelial cell (AT2)-associated signal and increased macrophage-associated signal during the acute phase (S2 Fig).

Because the gCap-associated signature changed early after elastase exposure, we next examined the expression of the apelin-APJ pathway. Lung *Apln* mRNA and apelin protein levels were reduced at day 1 after elastase exposure, and *Agtrl1*/APJ transcript expression also decreased during the acute phase (Fig 1F and 1G). Together, within this bulk transcriptomic framework, these findings suggest that elastase-induced injury is associated with an early reduction in gCap-associated transcriptional features and with decreased expression of the apelin-APJ pathway.

### Early apelin receptor activation preserves APJ-positive endothelial staining and reduces acute neutrophil-dominant inflammation after elastase exposure

We next tested whether early activation of the apelin receptor modifies endothelial-associated and inflammatory responses following elastase exposure. [Pyr^1^]-Apelin-13 was administered intraperitoneally beginning 1 hour after elastase exposure, and lungs and bronchoalveolar lavage fluid were analyzed at day 1 (Fig 2A). APJ/CD31 immunofluorescence showed that APJ-positive endothelial staining was reduced after elastase exposure and partially preserved by [Pyr^1^]-Apelin-13 treatment (S3 Fig). This finding provided a histological index of altered APJ-positive endothelial signal during the acute phase, complementing the transcriptomic reduction in apelin-APJ pathway expression observed after elastase injury.

**Fig 2.**
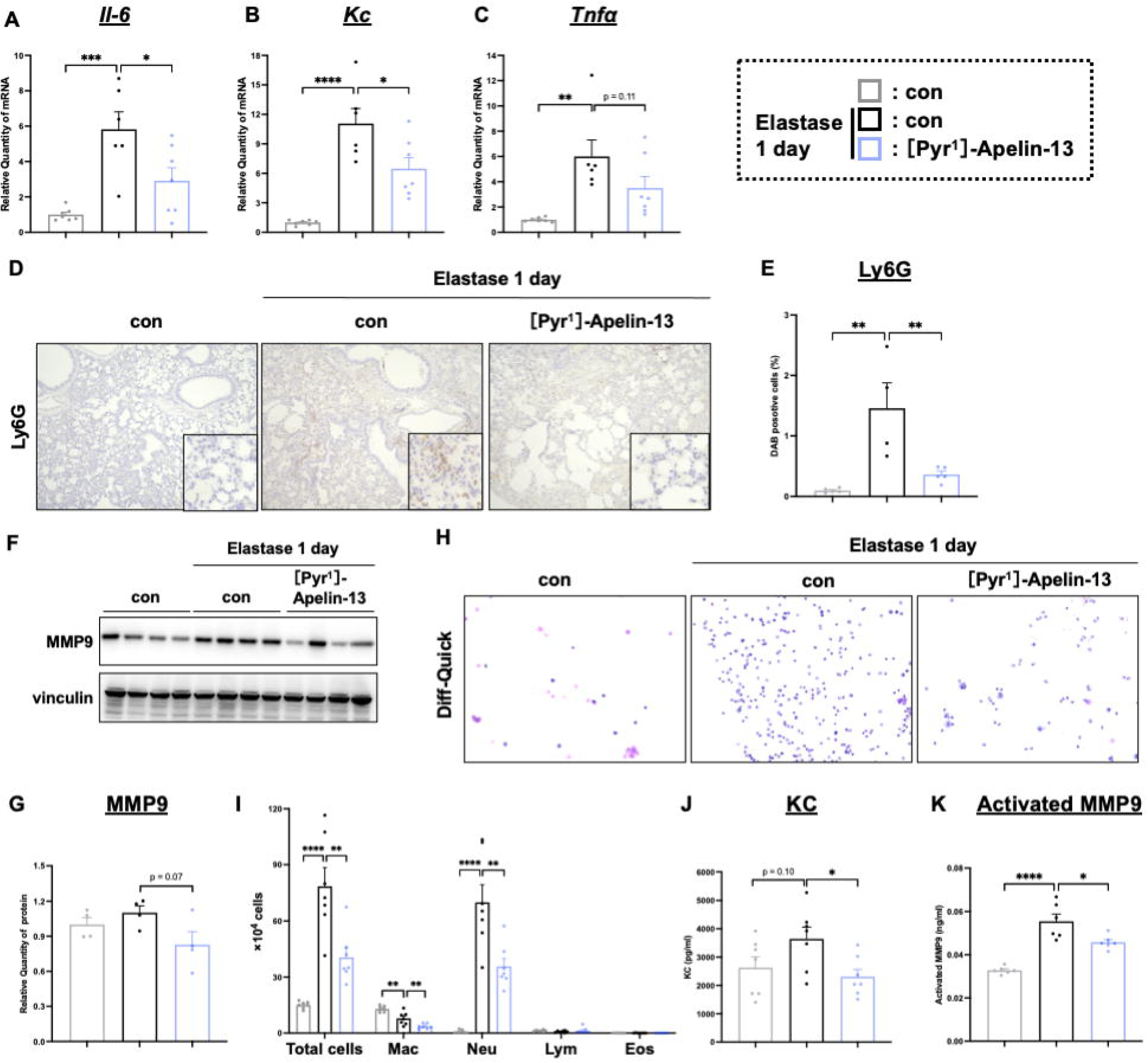
Early [Pyr^1^]-Apelin-13 treatment suppresses acute elastase-induced inflammation. (A) Experimental design for acute [Pyr^1^]-Apelin-13 administration after elastase exposure. (B–D) Relative lung mRNA expression of *Il6*, *Kc*/*Cxcl1*, and *Tnfα* at day 1. (E, F) Representative Ly6G immunohistochemistry and quantification of Ly6G-positive neutrophil accumulation in lung sections. (G, H) Lung MMP-9 protein levels assessed by immunoblotting and densitometry. (I) Total and differential bronchoalveolar lavage fluid cell counts at day 1. (J) Bronchoalveolar lavage fluid KC/CXCL1 concentration. (K) Bronchoalveolar lavage fluid matrix MMP-9 activity. Data are means ± SEM; n = 6–7 mice per group. *P* values were determined by one-way ANOVA with Dunnett’s multiple-comparisons test. \**P* < 0.05, \*\**P* < 0.01, \*\*\**P* < 0.001, \*\*\*\**P* < 0.0001.

Elastase exposure increased lung expression of *Il6*, *Kc*/*Cxcl1*, and *Tnfα*. [Pyr^1^]-Apelin-13 treatment reduced *Il6* and *Kc*/*Cxcl1* expression and showed a suppressive trend for *Tnfα* (Fig 2B–2D). Immunohistochemistry showed that Ly6G-positive neutrophil accumulation was increased after elastase exposure and was reduced by [Pyr^1^]-Apelin-13 treatment (Fig 2E and 2F). Bronchoalveolar lavage fluid (BALF) analysis also showed that elastase-induced increases in total cells and neutrophils were reduced by [Pyr^1^]-Apelin-13 (Fig 2I).

Matrix metalloproteinase-9 (MMP-9) protein expression in lung tissue tended to decrease with [Pyr^1^]-Apelin-13 treatment, and MMP-9 activity in BALF was significantly reduced (Fig 2G, 2H, and 2K). KC/CXCL1 levels in BALF were also reduced by [Pyr^1^]-Apelin-13 treatment (Fig 2J). Lung *Mmp9* mRNA expression was suppressed, whereas Iba1-positive cell accumulation and *Mmp12* expression were less clearly affected (S4 Fig). Together, these findings indicate that early activation of the apelin receptor preserves APJ-positive endothelial staining, attenuates acute neutrophil-dominant inflammation, and reduces MMP-9 activity following elastase injury. The parallel decreases in inflammatory cytokine expression, Ly6G-positive neutrophil accumulation, BALF neutrophilia, and MMP-9 activity support broad suppression of the acute inflammatory response.

### Early, but not delayed, apelin receptor activation attenuates elastase-induced airspace enlargement

We then assessed whether early suppression of acute inflammation by apelin receptor activation was associated with reduced emphysematous remodeling. [Pyr^1^]-Apelin-13 was administered once daily beginning 1 hour after elastase exposure and continued for 3 weeks (Fig 3A). Pulmonary function was evaluated at 3 weeks. Elastase exposure altered pulmonary function parameters consistent with emphysema development. Early [Pyr^1^]-Apelin-13 treatment did not significantly improve inspiratory capacity (IC), compliance, elastance, forced expiratory volume in 0.1 second (FEV_0.1_), or forced vital capacity (FVC), although the FEV_0.1_/FVC ratio showed a trend toward improvement (Fig 3B–3G). Histological analysis showed marked elastase-induced airspace enlargement, whereas mean linear intercept (MLI) was significantly reduced in [Pyr^1^]-Apelin-13-treated mice (Fig 3H and 3I).

**Fig 3.**
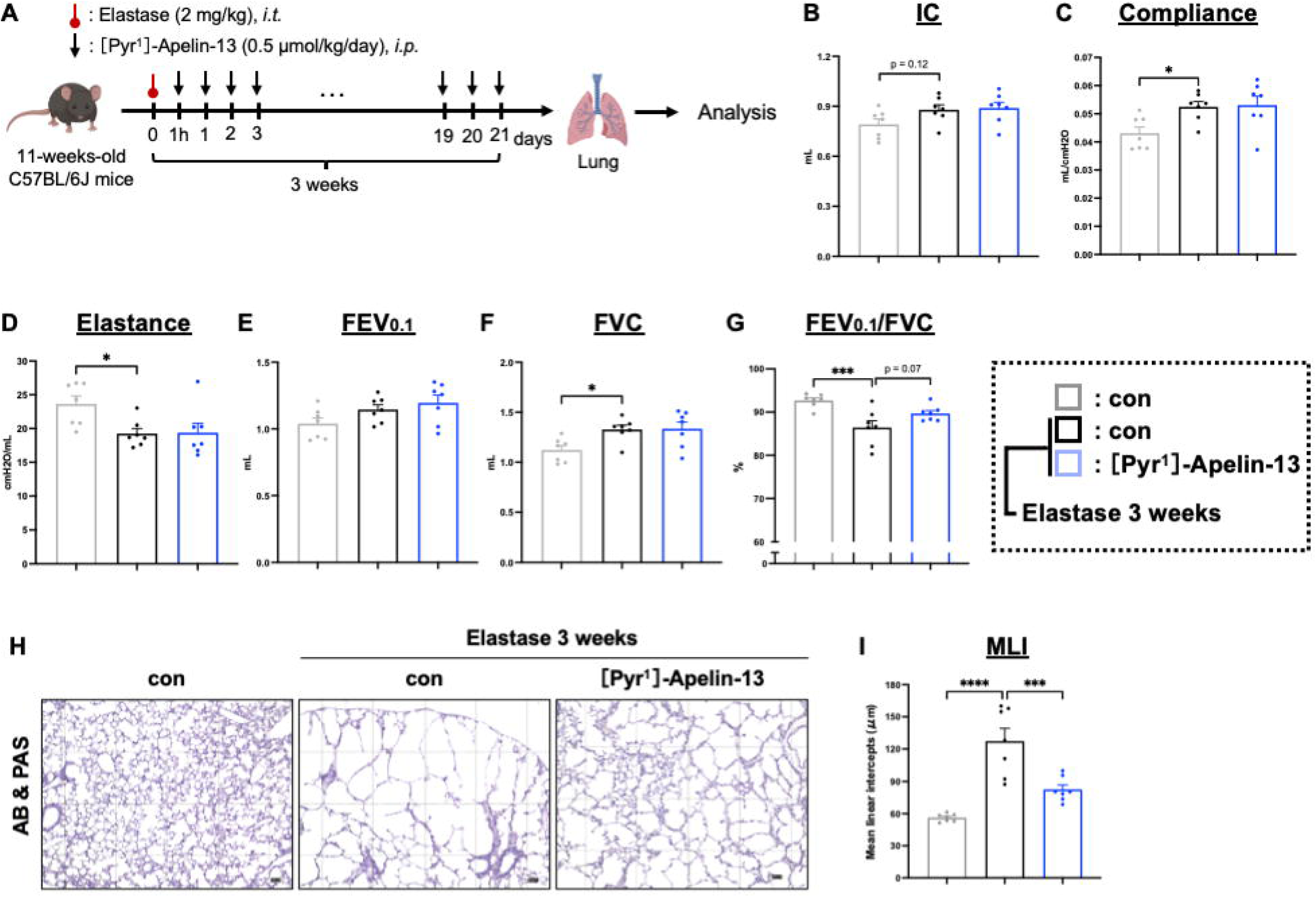
Early administration of [Pyr^1^]-Apelin-13 attenuates elastase-induced airspace enlargement. (A) Experimental design for early [Pyr^1^]-Apelin-13 treatment. Treatment was initiated 1 hour after elastase exposure and continued once daily for 3 weeks. (B–G) Pulmonary function parameters, including inspiratory capacity (IC), compliance, elastance, forced expiratory volume in 0.1 second (FEV_0.1_), forced vital capacity (FVC), and FEV_0.1_/FVC ratio. (H, I) Representative Alcian blue/periodic acid-Schiff-stained lung sections and quantification of mean linear intercept. Data are means ± SEM; n = 6–7 mice per group. *P* values were determined by one-way ANOVA with Dunnett’s multiple-comparisons test. \**P* < 0.05, \*\*\**P* < 0.001, \*\*\*\**P* < 0.0001.

To determine whether treatment timing was important, [Pyr^1^]-Apelin-13 administration was initiated 3 weeks after elastase exposure, when emphysema was established. Delayed treatment did not improve pulmonary function or MLI (S5 Fig). These findings suggest that apelin receptor activation is more effective during the acute injury phase than after emphysematous remodeling has already developed.

### Chronic **β**ENaC-transgenic airway disease shows a different inferred endothelial subtype pattern and does not respond to apelin receptor activation

The elastase model showed an early reduction in a gCap-associated signature and in apelin pathway expression. We therefore examined whether a similar pattern was present in C57BL/6J-βENaC-transgenic mice, which develop chronic airway inflammation, mucus obstruction, and obstructive lung disease-like phenotypes [24]. Bulk RNA-sequencing data from βENaC-transgenic mice were analyzed by CIBERSORTx using the same mouse lung single-cell reference dataset (Fig 4A).

**Fig 4.**
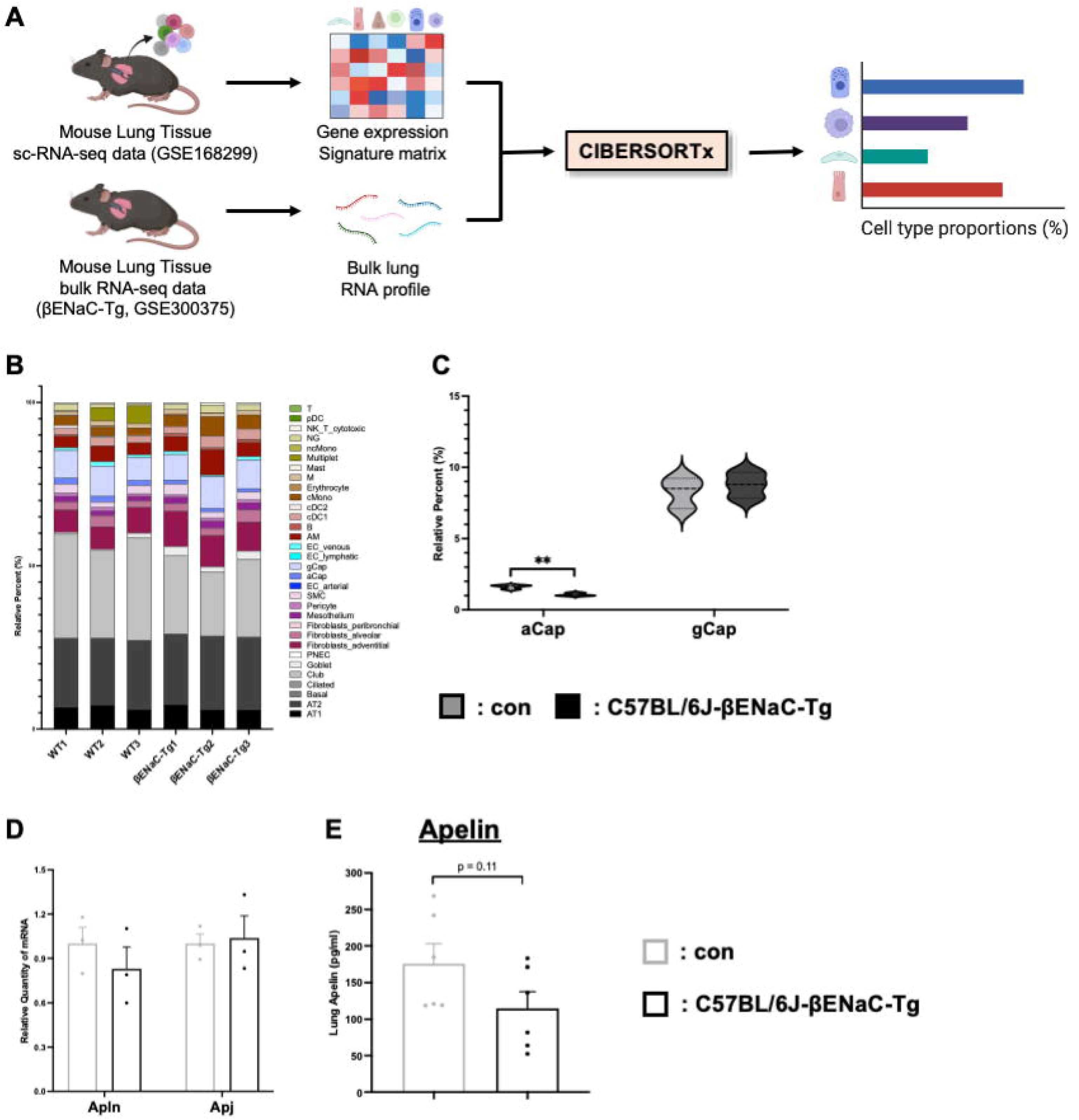
βENaC-transgenic mice show a distinct inferred endothelial subtype pattern and limited response to apelin receptor activation. (A) Overview of CIBERSORTx analysis using a mouse lung single-cell RNA-sequencing reference dataset and bulk lung RNA-sequencing data from control and C57BL/6J-βENaC-transgenic mice. (B) Estimated proportions of lung cell-type-associated signatures inferred by CIBERSORTx. (C) Inferred aCap and gCap fractions. (D) Lung expression of *Apln* and apelin receptor/APJ-associated transcript in bulk RNA-sequencing data. (E) Lung apelin protein levels measured by enzyme-linked immunosorbent assay. Data are means ± SEM; n = 3–6 mice per group. *P* values were determined by two-tailed Student’s *t*-test. \*\**P* < 0.01.

In contrast to the elastase model, the inferred gCap fraction was not reduced in βENaC-transgenic mice, whereas the inferred aCap fraction was decreased (Fig 4B and 4C). Other epithelial, mesenchymal, and immune cell-associated changes were also detected (S6 Fig). *Apln* and *Agtrl1*/APJ-associated transcript expression showed only modest changes, and lung apelin protein levels were modestly reduced without reaching statistical significance (Fig 4D and 4E). Consistent with these differences, intraperitoneal [Pyr^1^]-Apelin-13 treatment did not improve pulmonary function or MLI in βENaC-transgenic mice (S7 Fig). These findings indicate that the endothelial subtype-associated transcriptional pattern and pharmacologic response to apelin receptor activation differ between acute elastase-induced injury and chronic βENaC-associated airway disease.

## Discussion

In this study, we investigated endothelium-associated transcriptional changes following elastase-induced lung injury and tested whether early activation of the apelin receptor modifies inflammation and emphysema development in mice. Bulk RNA sequencing with CIBERSORTx-based deconvolution suggested an early reduction in a gCap-associated signature following elastase exposure, accompanied by decreased expression of the apelin-apelin receptor pathway. Pharmacologic activation of the apelin receptor with [Pyr^1^]-Apelin-13 reduced acute inflammatory mediator expression, Ly6G-positive neutrophil accumulation, bronchoalveolar lavage neutrophilia, and MMP-9 activity. Early treatment attenuated subsequent airspace enlargement, whereas delayed treatment after emphysema was established did not improve disease outcomes.

A key finding of this study is the timing-dependent effect of apelin receptor activation. Early [Pyr^1^]-Apelin-13 treatment reduced acute neutrophilic inflammation and later histological emphysema, whereas delayed treatment did not reverse established emphysematous remodeling. These results suggest that apelin receptor activation primarily modulates early injury and inflammatory responses rather than repairing established structural destruction. The effect on lung function was modest compared with the effect on MLI, indicating that the histological benefit did not translate into robust physiological improvement under the conditions tested.

The endothelial interpretation of these findings requires caution. CIBERSORTx is useful for estimating cell-type-associated composition from bulk RNA-sequencing data, but its results depend on the reference dataset, normalization strategy, and the stability of cell-type-specific transcriptional signatures [26]. Inflammatory injury may also alter transcriptional states within individual cell populations, thereby influencing deconvolution estimates. Thus, the observed reduction in the inferred gCap fraction is best interpreted as a decrease in gCap-associated transcriptional features rather than as direct evidence of reduced gCap abundance. Consistent with this interpretation, APJ/CD31 immunofluorescence supported reduced apelin receptor-positive endothelial staining after elastase exposure and its attenuation by [Pyr^1^]-Apelin-13 treatment, but APJ/CD31 staining alone does not define endothelial subtype identity.

The inflammatory data support an acute anti-inflammatory effect of [Pyr^1^]-Apelin-13 after elastase injury. Elastase-induced *Il6*, *Kc*/*Cxcl1*, and *Tnfα* expression was reduced, Ly6G-positive neutrophil accumulation was suppressed, and bronchoalveolar lavage neutrophilia decreased. MMP-9 activity was also reduced. Neutrophils are important contributors to protease-mediated extracellular matrix degradation in emphysema [29,30], and the parallel reduction in neutrophilic inflammation and MMP-9 activity provides a plausible link to the attenuation of airspace enlargement. Because MMP-9 and inflammatory mediators can be produced by multiple lung cell types, these findings indicate suppression of a neutrophil-associated inflammatory milieu rather than defining the precise cellular source of each mediator.

The mechanisms by which apelin receptor activation modifies endothelial-associated responses remain to be determined. In other systems, apelin receptor signaling regulates endothelial survival, angiogenesis, barrier function, and vascular repair through pathways involving PI3K-Akt, ERK, and endothelial nitric oxide synthase [31–33]. The present study did not directly measure endothelial apoptosis, proliferation, or downstream apelin receptor signaling. Therefore, whether [Pyr^1^]-Apelin-13 acts through endothelial survival, proliferation, barrier stabilization, indirect immune modulation, or a combination of these mechanisms remains unresolved.

The comparison with βENaC-transgenic mice suggests that the effect of apelin receptor activation is context dependent. In the elastase model, early injury was associated with a reduced inferred gCap signature and decreased apelin pathway expression. In βENaC-transgenic mice, the inferred gCap fraction was maintained, the inferred aCap fraction was reduced, and apelin receptor activation did not improve pulmonary function or histological emphysema-like changes. These findings suggest that apelin receptor activation may be more relevant to acute distal injury-associated inflammatory responses than to chronic proximal airway-dominant disease. However, this comparison remains exploratory because the two models differ in initiating injury, disease kinetics, inflammatory environment, and remodeling pattern.

This study has several limitations. Human lung tissues were not analyzed, and no direct human validation was performed. Although prior human single-cell studies of COPD support the relevance of endothelial niche alterations [14], the present findings require validation in human samples stratified by disease stage and inflammatory phenotype. In addition, endothelial subtype composition was inferred from bulk transcriptomes and was not directly validated by flow cytometry or definitive multi-marker histology. APJ/CD31 staining provided supportive information on apelin receptor-positive endothelial signals but did not specifically identify gCap. Finally, apoptosis and proliferation markers, downstream apelin receptor signaling, and public human or cigarette smoke-associated single-cell datasets were not analyzed. Future studies integrating elastase, cigarette smoke, and human COPD datasets, with direct validation of endothelial subtypes, will be needed to determine whether apelin receptor-associated endothelial programs are conserved across models and species.

In conclusion, early activation of the apelin receptor reduced acute neutrophil-dominant inflammation and attenuated elastase-induced airspace enlargement in mice. These effects were associated with changes in endothelial apelin receptor-positive signals and inferred gCap-associated transcriptional features. The data support a time-sensitive protective effect of apelin receptor activation in the development of acute elastase-induced emphysema, while highlighting the need for direct validation of endothelial subtypes and mechanistic studies.

## Conclusions

Early activation of the apelin receptor with [Pyr^1^]-Apelin-13 reduced acute neutrophilic inflammation and attenuated airspace enlargement in elastase-induced emphysema. These findings support a time-sensitive role for apelin-APJ signaling in early injury responses.

## Supporting information

S1 Raw Images

## Acknowledgments

The authors have no additional acknowledgments.

## Supporting information captions

**S1 Fig. Bulk RNA-sequencing analysis of lung tissue in an elastase-induced emphysema mouse model.**

(A) Volcano plot of differentially expressed genes between control and elastase-treated mice at day 1.

(B) Gene Ontology enrichment analysis of differentially expressed genes in (A).

(C) Network plot of significantly enriched Gene Ontology (GO) terms among upregulated genes from (B).

(D) Volcano plot comparing control and elastase-treated mice at week 1.

(E) GO enrichment analysis corresponding to (D).

(F) GO term-gene network of upregulated biological processes from (E).

(G) Volcano plot comparing control and elastase-treated mice at week 2.

(H) GO enrichment analysis corresponding to (G).

(I) GO network plot of enriched biological processes among upregulated genes from (H).

Analyses were performed using RNAseqChef. Data are means ± SEM; n = 3 mice per group.

**S2 Fig. Predicted cellular composition in lungs from elastase-induced emphysema mice.**

(A–C) Violin plots of inferred cell-type fractions in control versus elastase-treated lungs based on the CIBERSORTx deconvolution shown in Fig 1D. (A) Epithelial cells, (B) mesenchymal cells, and (C) immune and other cell types.

Data are means ± SEM; n = 3 mice per group. *P* values were determined by one-way ANOVA with Dunnett’s multiple-comparisons test. \**P* < 0.05, \*\**P* < 0.01, \*\*\**P* < 0.001.

**S3 Fig. APJ/CD31 immunofluorescence after elastase injury and [Pyr^1^]-Apelin-13 treatment.**

(A) Experimental design. [Pyr^1^]-Apelin-13 was administered after elastase exposure, and lungs were analyzed at day 1.

(B, C) Representative immunofluorescence images and quantification of APJ/CD31-positive endothelial signal in lung sections from control, elastase-treated, and [Pyr^1^]-Apelin-13-treated elastase mice. Sections were stained for APJ, CD31, and DAPI. These data are presented as supportive evidence of apelin receptor-positive endothelial signal and not as definitive identification of general capillary endothelial cells.

Data are means ± SEM; n = 4–5 mice per group. *P* values were determined by one-way ANOVA with Dunnett’s multiple-comparisons test. \**P* < 0.05, \*\**P* < 0.01.

**S4 Fig. Effects of [Pyr^1^]-Apelin-13 on additional inflammatory markers after elastase exposure.**

(A) Relative *Mmp9* mRNA expression in lung tissue from control, elastase-treated, and [Pyr^1^]-Apelin-13-treated elastase mice at day 1.

(B, C) Representative Iba1 immunohistochemistry and quantification of Iba1-positive cell accumulation in lung sections at day 1.

(D) Relative *Mmp12* mRNA expression in lung tissue at day 1.

Data are means ± SEM; n = 6–7 mice per group. *P* values were determined by one-way ANOVA with Dunnett’s multiple-comparisons test. \**P* < 0.05, \*\**P* < 0.01, \*\*\**P* < 0.001.

**S5 Fig. Delayed [Pyr^1^]-Apelin-13 administration does not attenuate established elastase-induced emphysema.**

(A) Experimental design. [Pyr^1^]-Apelin-13 was initiated 3 weeks after elastase exposure and continued through week 4.

(B–G) Pulmonary function parameters measured using the flexiVent system. (H, I) Morphometric assessment of emphysema severity by MLI.

Data are means ± SEM; n = 6–7 mice per group. *P* values were determined by one-way ANOVA with Dunnett’s multiple-comparisons test.

**S6 Fig. Predicted cellular composition in lungs from C57BL/6J-βENaC-transgenic mice.**

(A–C) Violin plots of inferred cell-type fractions in control versus C57BL/6J-βENaC-transgenic lungs based on the CIBERSORTx deconvolution shown in Fig 4B. (A) Epithelial cells, (B) mesenchymal cells, and (C) immune and other cell types.

Data are means ± SEM; n = 3 mice per group. *P* values were determined by two-tailed Student’s *t*-test. \**P* < 0.05.

**S7 Fig. Effects of intraperitoneal [Pyr^1^]-Apelin-13 on COPD-like phenotypes in C57BL/6J-βENaC-transgenic mice.**

(A) Experimental design for [Pyr^1^]-Apelin-13 administration in C57BL/6J-βENaC-transgenic mice.

(B–G) Pulmonary function parameters measured using the flexiVent system in control, C57BL/6J-βENaC-transgenic, and [Pyr^1^]-Apelin-13-treated C57BL/6J-βENaC-transgenic mice.

(H, I) Morphometric assessment by mean linear intercept.

Data are means ± SEM; n = 5–6 mice per group. *P* values were determined by one-way ANOVA with Dunnett’s multiple-comparisons test. \*\**P* < 0.01, \*\*\**P* < 0.001, \*\*\*\**P* < 0.0001.

## Data Availability Statement

Bulk RNA-sequencing data from the elastase-induced emphysema model are available in GEO under accession number GSE324765. Bulk RNA-sequencing data from C57BL/6J-βENaC-transgenic mice are available in GEO under accession number GSE300375. The raw data supporting the conclusions of this article will be made available by the authors. Original uncropped immunoblot images are provided in S1 Raw Images. The mouse lung single-cell RNA-sequencing reference dataset used for CIBERSORTx analysis is publicly available under accession number GSE168299.

## Author contributions

Conceptualization: T.K., R.N., H.K., T.S.

Data curation: T.K., T.S.

Formal analysis: T.K., T.S.

Funding acquisition: T.S.

Investigation: T.K., K.K., M.U., K.N., N.T., C.O.

Methodology: T.K., R.N., T.S.

Project administration: H.K., T.S.

Supervision: H.K., T.S.

Writing – original draft: T.K., M.A.S., T.S.

Writing – review & editing: T.K., M.A.S., T.S.

