## Supplementary material for "Early apelin receptor activation attenuates elastase-induced emphysema and preserves endothelial apelin receptor signaling in mice": S1 Raw Images: S1 Raw Images.pdf

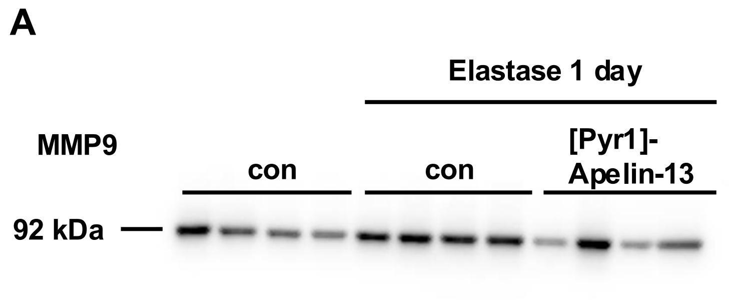


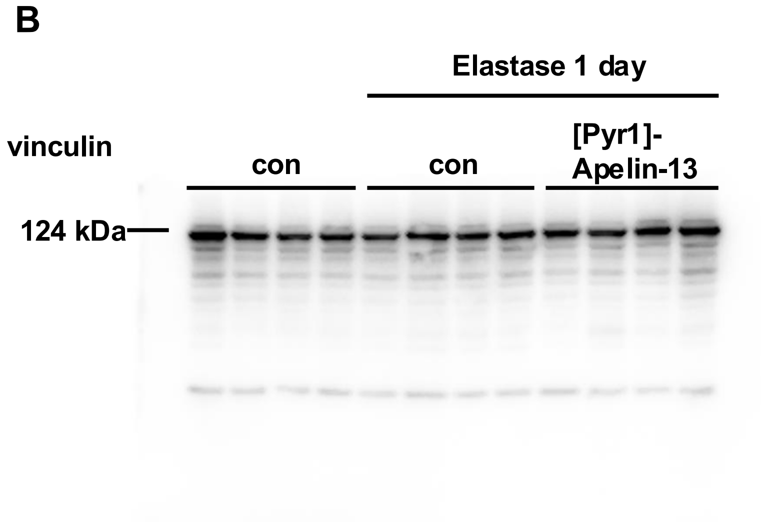


A-B: The relative quantity of protein levels of MMP9 in the lung tissue of WT, elastase-induced emphysema model, and [$Pyr^1$]-Apelin-13-treated emphysema model mice at 1 day after elastase administration by western blotting.
