## Supplementary material for "Early apelin receptor activation attenuates elastase-induced emphysema and preserves endothelial apelin receptor signaling in mice": S1-S7 Figures: S1-S7 Figures.pdf

### S1-S7 Figs

### S1 Fig

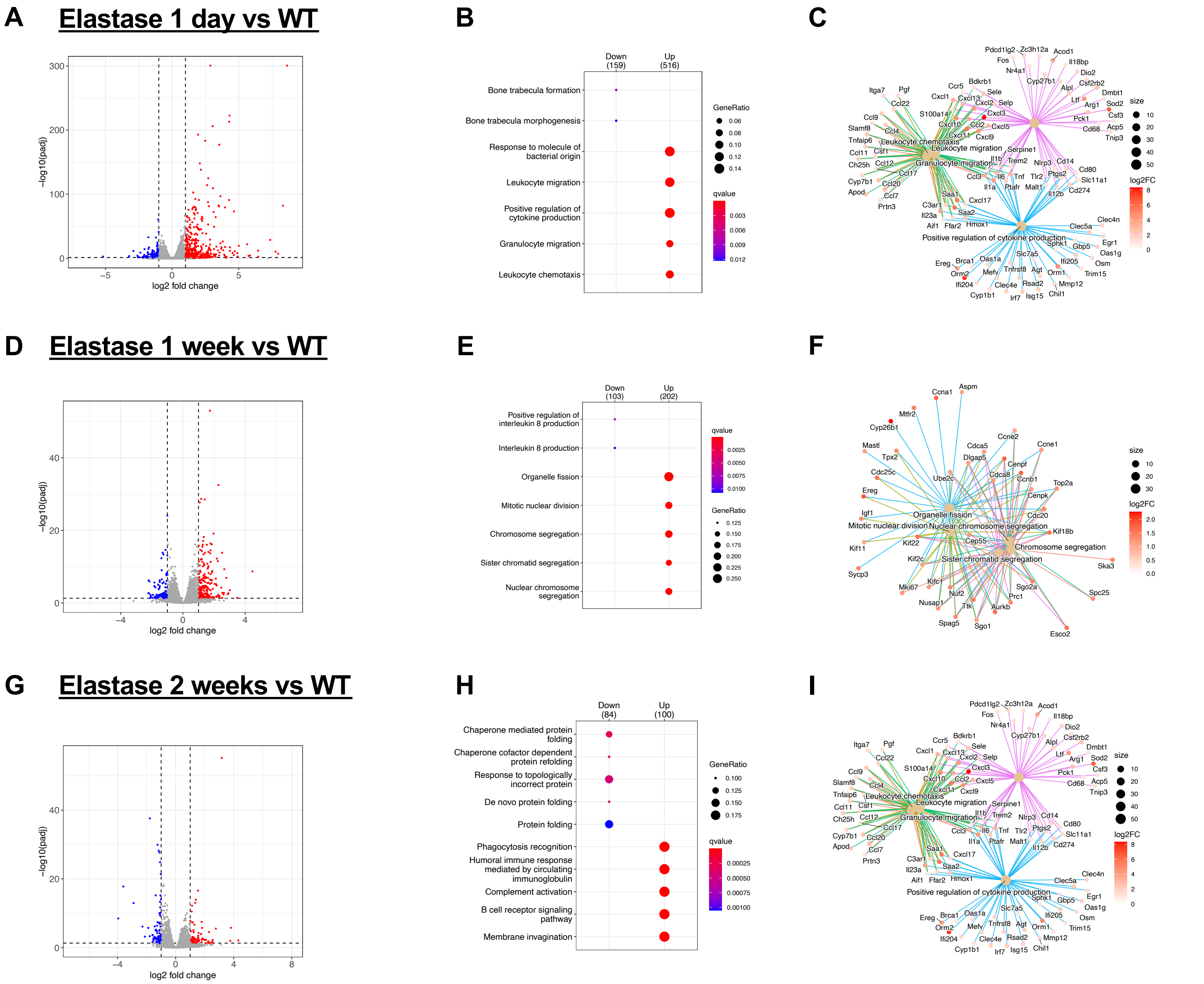


**S1 Fig. Bulk RNA-sequencing analysis of lung tissue in an elastase-induced emphysema mouse model.**

(A) Volcano plot of differentially expressed genes between control and elastase-treated mice at day 1.
(B) Gene Ontology enrichment analysis of differentially expressed genes in (A).
(C) Network plot of significantly enriched Gene Ontology (GO) terms among upregulated genes from (B).
(D) Volcano plot comparing control and elastase-treated mice at week 1.
(E) GO enrichment analysis corresponding to (D).
(F) GO term-gene network of upregulated biological processes from (E).
(G) Volcano plot comparing control and elastase-treated mice at week 2.
(H) GO enrichment analysis corresponding to (G).
(I) GO network plot of enriched biological processes among upregulated genes from (H).
Analyses were performed using RNAseqChef.
Data are means ± SEM; n = 3 mice per group.

### S2 Fig

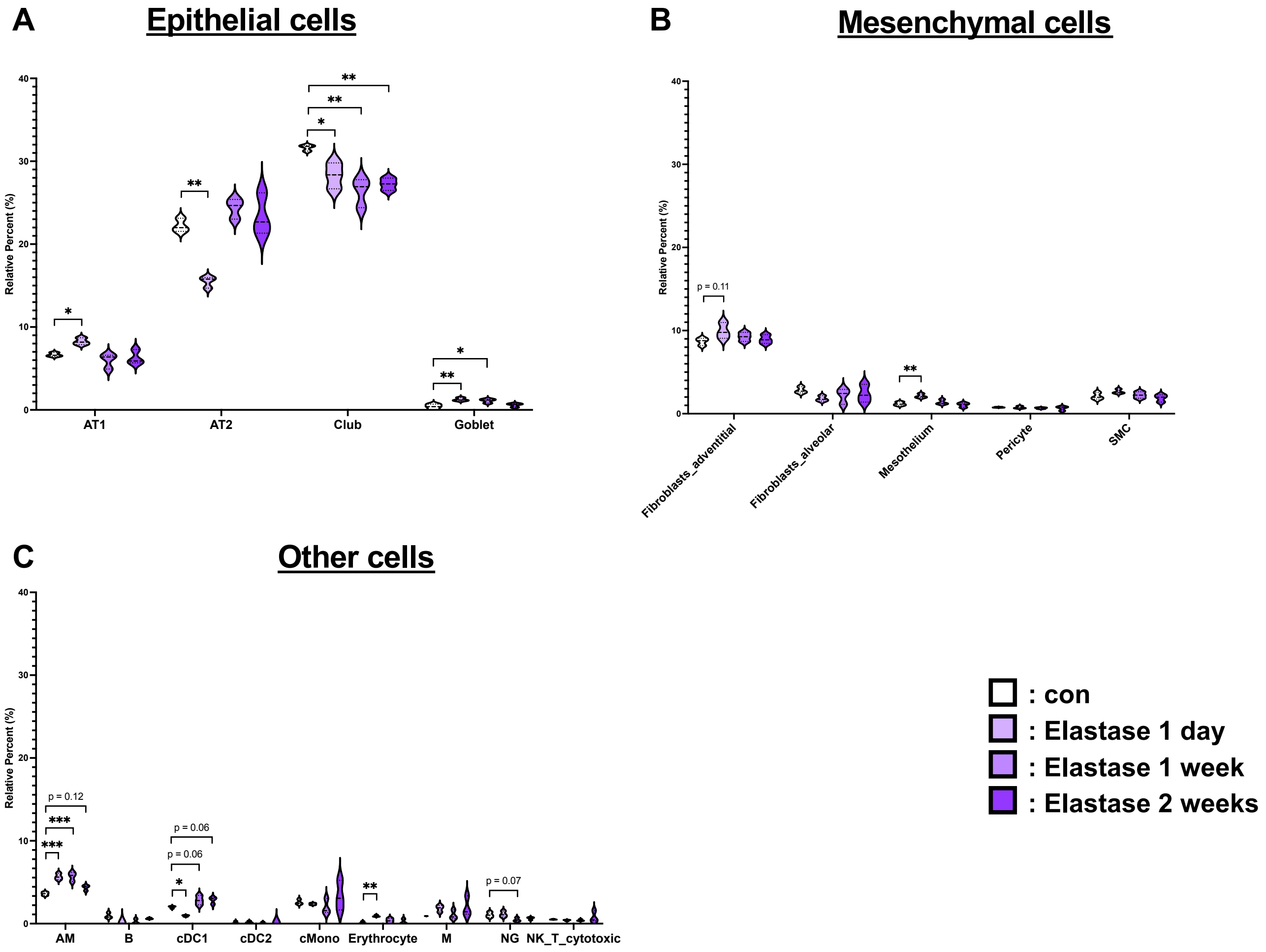


**S2 Fig. Predicted cellular composition in lungs from elastase-induced emphysema mice.**

(A–C) Violin plots of inferred cell-type fractions in control versus elastase-treated lungs based on the CIBERSORTx deconvolution shown in Fig 1D. (A) Epithelial cells, (B) mesenchymal cells, and (C) immune and other cell types.

Data are means ± SEM; n = 3 mice per group. P values were determined by one-way ANOVA with Dunnett's multiple-comparisons test. *$P < 0.05$, **$P < 0.01$, ***$P < 0.001$.

#### S3 Fig

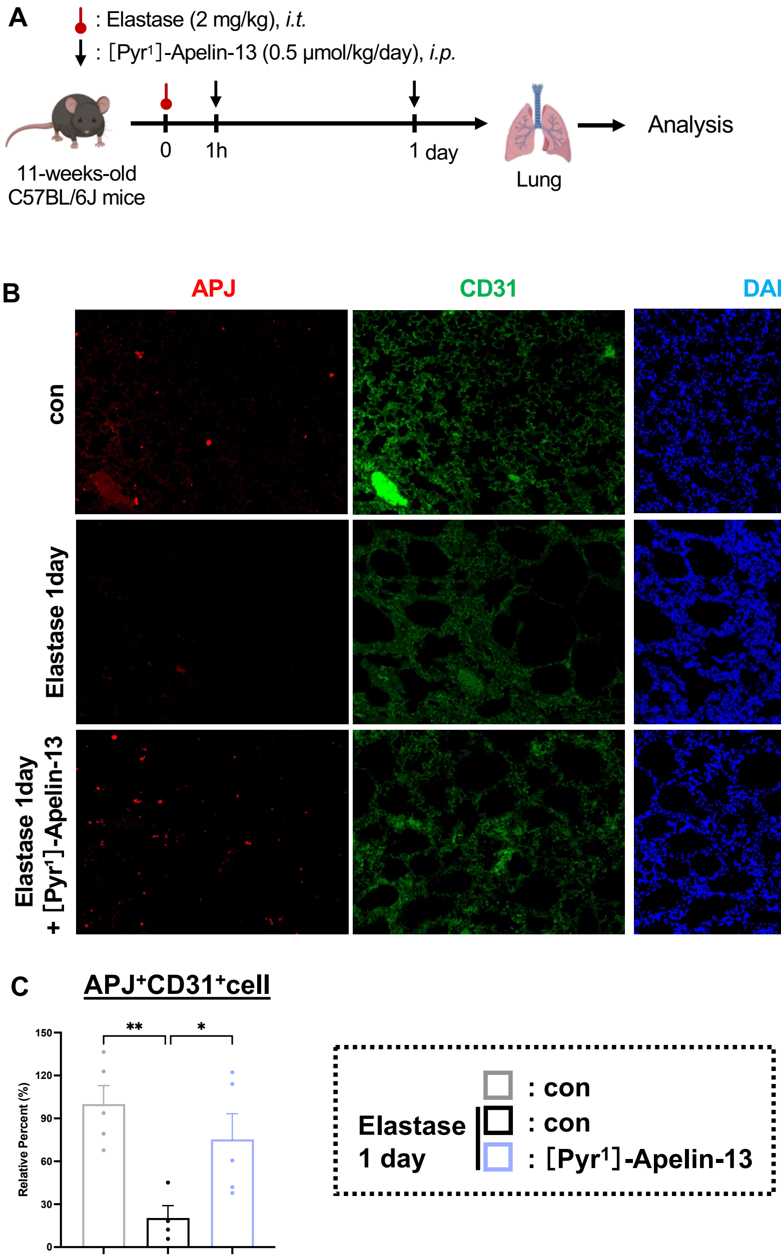


**S3 Fig. APJ/CD31 immunofluorescence after elastase injury and [Pyr[1]]-Apelin-13 treatment.**

(A) Experimental design. [Pyr[1]]-Apelin-13 was administered after elastase exposure, and lungs were analyzed at day 1.

(B, C) Representative immunofluorescence images and quantification of APJ/CD31-positive endothelial signal in lung sections from control, elastase-treated, and [Pyr[1]]-Apelin-13-treated elastase mice. Sections were stained for APJ, CD31, and DAPI. These data are presented as supportive evidence of apelin receptor-positive endothelial signal and not as definitive identification of general capillary endothelial cells.

Data are means ± SEM; n = 4–5 mice per group. *P* values were determined by one-way ANOVA with Dunnett's multiple-comparisons test. $*P < 0.05$, $**P < 0.01$.

### S4 Fig

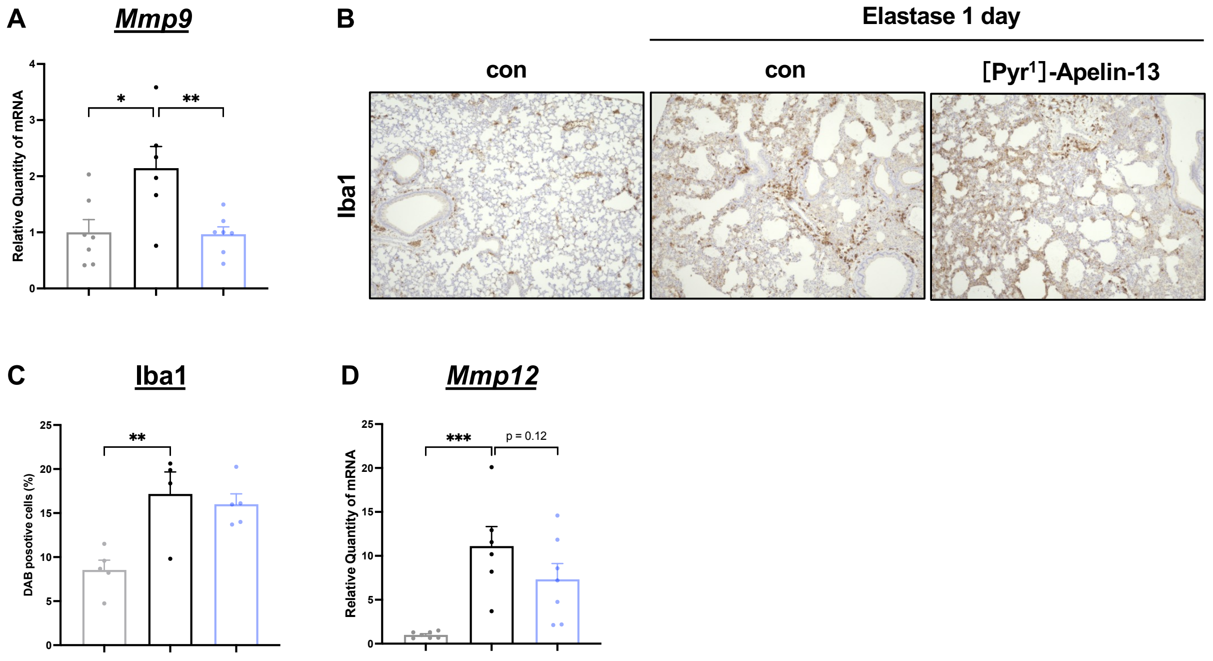


**S4 Fig. Effects of [Pyr1]-Apelin-13 on additional inflammatory markers after elastase exposure.**

(A) Relative *Mmp9* mRNA expression in lung tissue from control, elastase-treated, and [Pyr1]-Apelin-13-treated elastase mice at day 1.

(B, C) Representative Iba1 immunohistochemistry and quantification of Iba1-positive cell accumulation in lung sections at day 1.

(D) Relative *Mmp12* mRNA expression in lung tissue at day 1.

Data are means ± SEM; n = 6–7 mice per group. *P* values were determined by one-way ANOVA with Dunnett's multiple-comparisons test. $^{*}P < 0.05$, $^{**}P < 0.01$, $^{***}P < 0.001$.

### S5 Fig

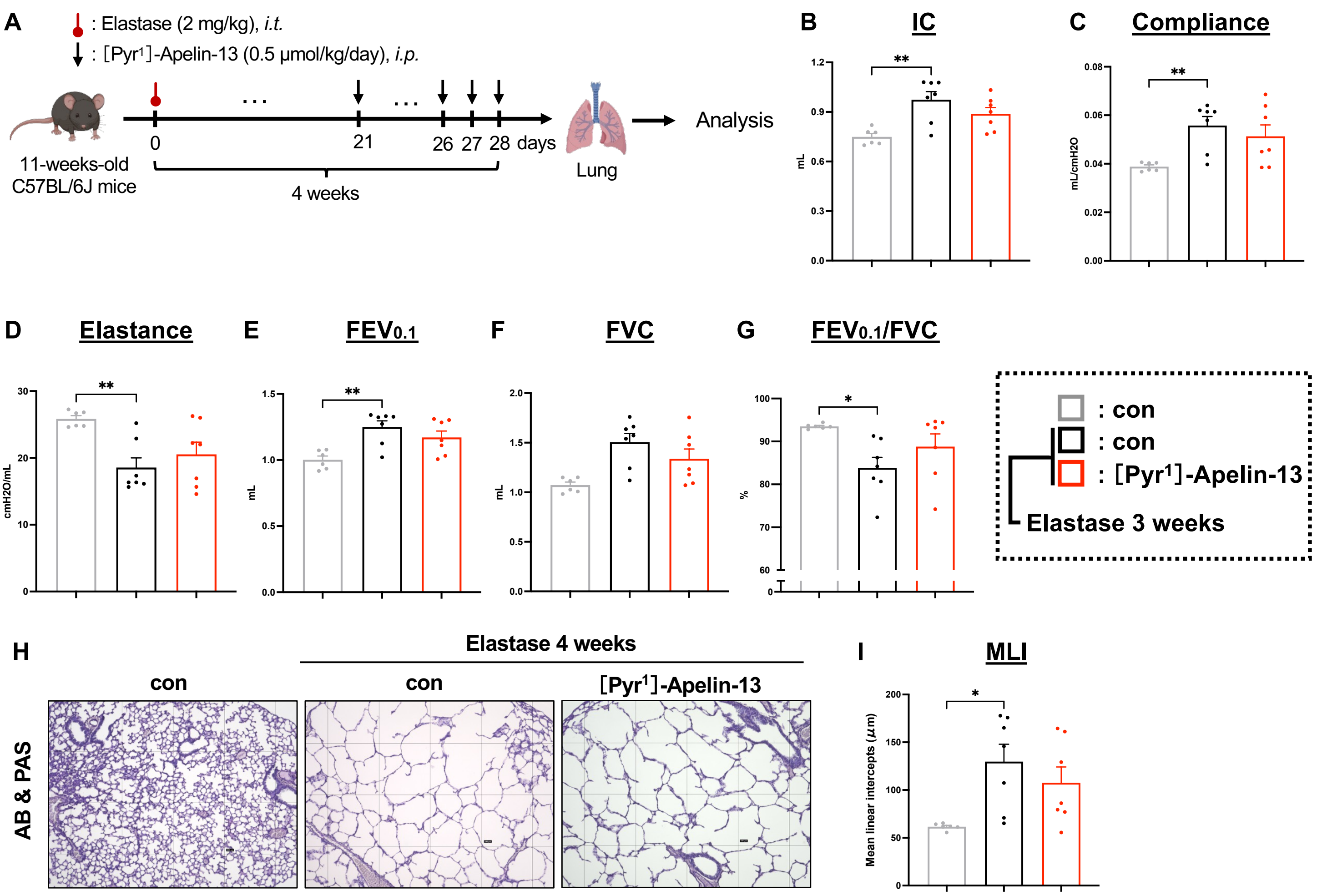


**S5 Fig. Delayed [Pyr¹]-Apelin-13 administration does not attenuate established elastase-induced emphysema.**

(A) Experimental design. [Pyr¹]-Apelin-13 was initiated 3 weeks after elastase exposure and continued through week 4.

(B–G) Pulmonary function parameters measured using the flexiVent system.

(H, I) Morphometric assessment of emphysema severity by MLI.

Data are means ± SEM; n = 6–7 mice per group. *P* values were determined by one-way ANOVA with Dunnett's multiple-comparisons test.

### S6 Fig

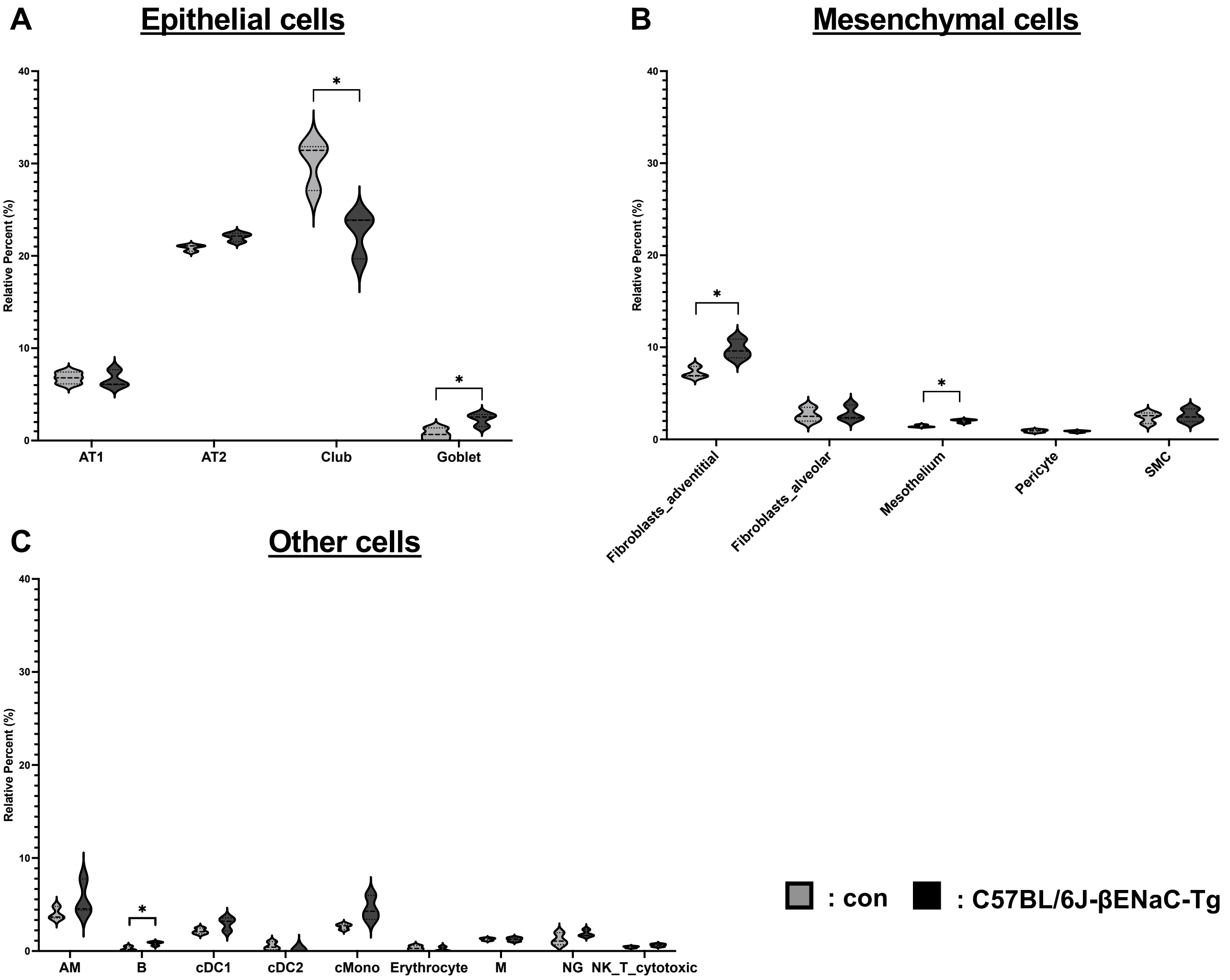


**S6 Fig. Predicted cellular composition in lungs from C57BL/6J-βENaC-transgenic mice.** (A–C) Violin plots of inferred cell-type fractions in control versus C57BL/6J-βENaC-transgenic lungs based on the CIBERSORTx deconvolution shown in Fig 4B. (A) Epithelial cells, (B) mesenchymal cells, and (C) immune and other cell types.

Data are means ± SEM; n = 3 mice per group. *P* values were determined by two-tailed Student's *t*-test. *$P < 0.05$.

### S7 Fig

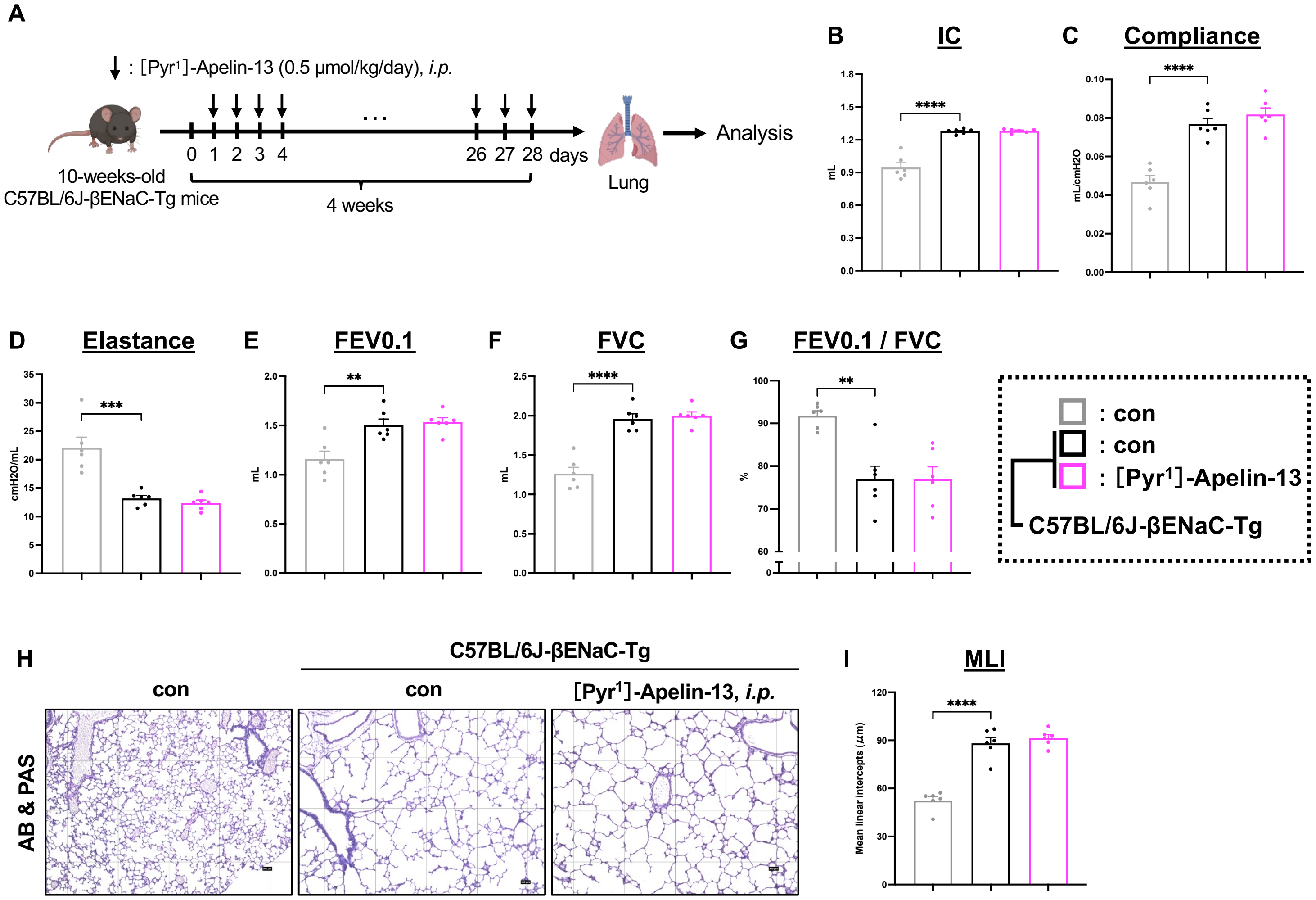


**S7 Fig. Effects of intraperitoneal [Pyr¹]-Apelin-13 on COPD-like phenotypes in C57BL/6J-βENaC-transgenic mice.**

(A) Experimental design for [Pyr¹]-Apelin-13 administration in C57BL/6J-βENaC-transgenic mice.

(B–G) Pulmonary function parameters measured using the flexiVent system in control, C57BL/6J-βENaC-transgenic, and [Pyr¹]-Apelin-13-treated C57BL/6J-βENaC-transgenic mice.

(H, I) Morphometric assessment by mean linear intercept.

Data are means ± SEM; n = 5–6 mice per group. *P* values were determined by one-way ANOVA with Dunnett's multiple-comparisons test. ***P* < 0.01, ****P* < 0.001, *****P* < 0.0001.
